# ModDotPlot—Rapid and interactive visualization of complex repeats

**DOI:** 10.1101/2024.04.15.589623

**Authors:** Alexander P. Sweeten, Michael C. Schatz, Adam M. Phillippy

**Affiliations:** Department of Computer Science, Johns Hopkins University, Baltimore, MD 21211, USA; Genome Informatics Section, Center for Genomics and Data Science Research, National Human Genome Research Institute, National Institutes of Health, Bethesda, MD 20892, USA

## Abstract

**Motivation:** A common method for analyzing genomic repeats is to produce a sequence similarity matrix visualized via a dot plot. Innovative approaches such as StainedGlass have improved upon this classic visualization by rendering dot plots as a heatmap of sequence identity, enabling researchers to better visualize multi-megabase tandem repeat arrays within centromeres and other heterochromatic regions of the genome. However, computing the similarity estimates for heatmaps requires high computational overhead and can suffer from decreasing accuracy.

**Results:** In this work we introduce ModDotPlot, an interactive and alignment-free dot plot viewer. By approximating average nucleotide identity via a *k*-mer-based containment index, ModDotPlot produces accurate plots orders of magnitude faster than StainedGlass. We accomplish this through the use of a hierarchical modimizer scheme that can visualize the full 128 Mbp genome of *Arabidopsis thaliana* in under 5 minutes on a laptop. ModDotPlot is bundled with a graphical user interface supporting real-time interactive navigation of entire chromosomes.

**Availability and Implementation:** ModDotPlot is available at https://github.com/marbl/ModDotPlot.

## Introduction

Large tandemly repeating blocks of DNA, such as satellite repeats and their complex higher-order structures, are ubiquitous in many eukaryotic genomes, yet have been notoriously difficult to sequence and assemble. These motifs occur disproportionately in telomeric, centromeric, and heterochromatic regions of the genome (17), and are commonly referred to as genomic “dark matter” due to their prior absence from reference genomes (32). Recent advances in long-read sequencing and assembly tools have enabled genomics researchers to successfully assemble these complex regions, culminating in the first complete human genome (24) as well as important model organisms such as *Arabidopsis* (23) and non-human primates (20). More broadly, with tools such as Verkko (29) and hifiasm (UL) (6) now able to automatically assemble complete “telomere-to-telomere” chromosomes, developing new methods to analyze these previously dark regions of the genome has taken on new importance.

Traditionally, dot plots have been useful visualizations to characterize the structure of complex repeats (19). To generate such a plot, a sequence *S* is typically aligned with itself using software such as MUMmer (21), and plotted in a two dimensional space. This approach results in a set of line segments from [*x, y*] to [*x* + *l −* 1, *y* + *l −* 1] for all matches of length *l* (above some minimum length threshold) beginning at positions *x* and *y* in *S*. This yields a single diagonal line segment, representing the sequence aligned with itself, and all off-diagonal segments representing the location of paralogous repeat copies. If based on a gapped sequence alignment, these segments may also be colored by their average sequence identity, but the internal, fine-grained structure of the repeats cannot be represented by this technique.

To overcome this limitation, recent work by Vollger *et al*. introduced StainedGlass (33), which relies on a rasterized rather than vectorized approach. In this framework, the aim is to generate a similarity matrix *M*_*w*_ where each cell *M*_*w*_(*A*_*i*_, *B*_*j*_) relates two genomic intervals *A*_*i*_ and *B*_*j*_ of length *w* beginning at positions *wi* and *wj* in *S*. By re-framing the problem in terms of intervals rather than single bases, a percent identity can be computed between all pairs of intervals and the matrix *M*_*w*_ can be rendered as a heatmap where each cell (pixel) represents the percent identity between the two substrings at the corresponding interval positions. This technique has extended the previously binary dot plot into a rich spectrum of information and proven highly effective for visualizing patterns of sequence evolution within tandem repeat arrays of both humans and plants (18, 34).

Although heatmaps produced by StainedGlass have been useful in practice, the workflow used to generate them has inherent limitations. First, StainedGlass uses Minimap2 (16) to determine sequence identity by computing the number of matches, mismatches, insertions, and deletions between pairs of substrings. Minimap2’s alignment heuristic is not well-suited for repetitive sequences (31) and leads to long runtimes, especially for short tandem repeats. For example, a single 3 Mbp human centromere requires over one hour to plot when running on a high performance compute cluster.

Furthermore, StainedGlass partitions the input sequence into intervals of a fixed size. Similar substrings that are split across this boundary may fail to align, leading to inaccurate identity estimates.

To improve upon these limitations, we propose a *k*-mer-based approach that bypasses the computationally expensive requirement of sequence alignment. Estimating sequence identity from sets of *k*-length substrings (*k*-mers) has seen increasing use in genomics (25). Such tools typically utilize downsampling methods, such as minhash, to reduce the size of each *k*-mer set before estimating sequence identity using the Jaccard index or related set similarity measure.

In this work, we introduce ModDotPlot, a novel heatmap visualization tool that rapidly estimates sequence identity using hierarchical modimizers, a form of fractional minhashing (4, 11). Modimizers are defined as hashed *k*-mer values that have no remainder when divided by some number *s*, which we refer to as the sparsity. Here we restrict *s* to powers of two, *s* = 2^*d*^, which conveniently results in the set of modimizers being: (1) precisely those hash values with *d* zeros in their least significant bits, and (2) a strict subset of the modimizers defined by *s* = 2^*d*−1^. We use this efficient membership test and hierarchical property to efficiently downsample genomic *k*-mers at multiple levels of sparsity. We show that the resulting modimizers can be used to accurately estimate the average nucleotide identity (ANI) of two substrings, while being resistant to segmentation artifacts and orders of magnitude faster than StainedGlass. To conclude, we demonstrate ModDotPlot’s ability to elucidate the centromeric satellite structure of both plants and animals.

## Materials and Methods

ModDotPlot takes as input a list of sequences in FASTA format and outputs a self-identity heatmap for each sequence, as well as comparative heatmaps for all pairwise combinations of sequences. In describing our methods, we assume the construction of a self-identity heatmap, but the necessary modifications for constructing comparative heatmaps is straightforward. ModDotPlot can be run one of two ways, specified at runtime: *Static mode* produces a static image file for each plot, while *Interactive mode* builds a plot hierarchy using multiple modimizer values so that the plot resolution can be adjusted in real time as the user adjusts the zoom level. We outline the workflow of both possible modes of ModDotPlot in Figure 1.

**Figure 1.**
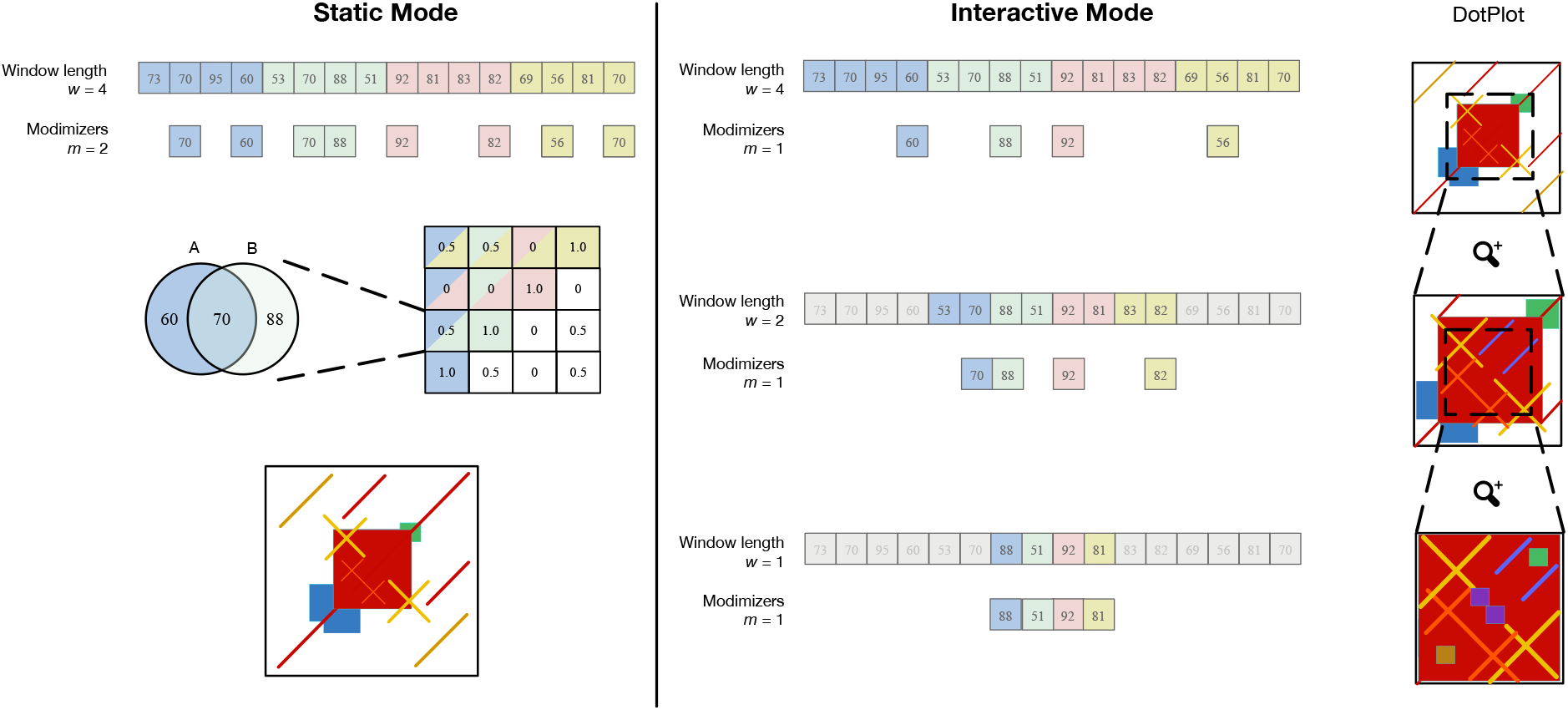
Overview of ModDotPlot’s workflow for producing a self-identity plot. **Static mode:** Hashed *k*-mers are evenly partitioned into intervals of length *w*. Modimizers are selected based on an estimated sketch size *m* within each interval. For each pairwise combination of intervals, identity is computed and stored in a matrix *M*_*w*_ . Finally, a heatmap is created based on the color thresholds provided. **Interactive mode:** Three distinct modimizer partitions are produced from a minimum interval length of *ŵ*=1 up to *w*=4. At launch, a heatmap is rendered for the largest window size (here, *w*=4). When the field of view is zoomed by half (highlighted region), the dot plot is rendered using a submatrix created from the partition at *w*=2. This process can extend until a plot produced from the minimum interval length *ŵ* is reached, with *m* remaining constant among all layers. While *m* = 1 is used here for demonstration, ModDotPlot adjusts the modimizer sparsity such that *m* ≈ 1000 in practice.

ModDotPlot first decomposes each sequence *S* of length *n* into a list of its constituent *k*-mers *S*_*k*_. Each *k*-mer and its reverse complement are passed through a hash function *h* : Ω *→* [0, *H*] for some *H* ∈ ℝ, with the smaller of the two values added into *S*_*k*_. Once broken down into *k*-mers, ModDotPlot partitions *S*_*k*_ into evenly sized and non-overlapping genomic intervals of size *w*, also referred to as the *window size*. We define the number of intervals as 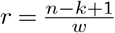, which we refer to as the *resolution*. This determines the height and width of the resulting heatmap. To reduce the runtime and space complexity of handling large sequences, ModDotPlot sketches each interval *A* into sets based on a modulo function, as originally proposed by Broder (4). We formally define our algorithm for sketching *S*_*k*_ in Supplementary Algorithm 1. This generates the following set for each interval:

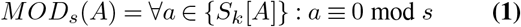

We refer to any *k*-mer present in the sketch *MOD*_*s*_(*A*) as a *modimizer*. We define *s ∈* ℤ^+^ as the *modimizer sparsity* and restrict *s* to powers of 2. Note that the sparsity value is inversely related to the number of modimizers selected (i.e. the density), with *s* = 2 resulting in approximately every second *k*-mer being selected, *s* = 4 with every fourth *k*-mer, and so on. Given a set of *k*-mers sampled from a long random string, the expected number of modimizers per window is:

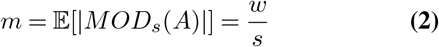

We refer to *m* as the *modimizer sketch size*, with larger values of *m* increasing the accuracy of the minhash similarity estimates. Given a desired plot resolution *r* and target sketch size *m*, the corresponding window size 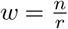 and required sparsity 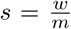 can be automatically derived. Based on prior work (25), we use *m* = 1, 000 as a good compromise between accuracy and efficiency.

In practice, if the *k*-mers in interval *A* are highly repetitive, then the true size of *MOD*_*s*_(*A*) can be significantly less than *m*. To avoid selecting too few *k*-mers in a window, we introduce a threshold set to half the expected number of modimizers. If the size of *MOD*_*s*_(*A*) is less than this threshold, modimizers are iteratively recomputed at half the sparsity until the modimizer count threshold is met or the sparsity hits one (i.e. every *k*-mer in *A* is included in the sketch).

Once the input sequence is partitioned and sketched, Mod-DotPlot produces a similiarity matrix *M*_*w*_ by estimating the identity between each pairwise combination of intervals *A* and *B*, which we refer to as a *cell* in the matrix. We estimate the proportion of *k*-mers in *A* that are contained in *B*, and vice-versa, via the containment index (4):

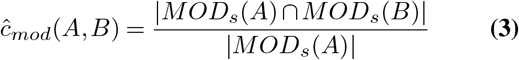

Hera *et al*. show that for the FracMinHash scheme, a correction factor is needed for an unbiased estimate of the containment index (28), to account for cases where |*MOD*_*s*_(*A*)| differs greatly from |*MOD*_*s*_(*B*)|. In practice, this can occur when interval *A* occurs in a repetitive genomic interval while interval *B* does not. Since modulo hashing is a variant of fractional minhashing, the same correction applies and we include the expected value in the denominator to achieve an unbiased estimate of the containment index:

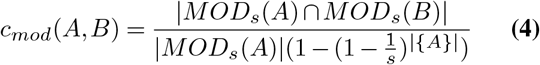

Furthermore, since the containment index drops exponentially with respect to the mutation rate (15), it is useful to represent this as an estimate of percent sequence identity. As implemented in MashScreen (26), we model the probability of mutation at each position in a *k*-mer with the binomial distribution to estimate the ANI as:

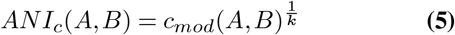

For self-identity plots, ModDotPlot sets *M*_*w*_(*A, B*) = max*{AN I*_*c*_(*A, B*), *ANI*_*c*_(*B, A*)*}* to ensure the resulting matrix is symmetric. We note that the containment index is not a distance metric, as it neither satisfies the symmetry property nor the triangle inequality property; however, for two equally sized intervals, we show that *ANI*_*c*_ correlates well with an alignment-based ANI. Furthermore, the containment index has the desirable property of not requiring a set operation in its denominator, meaning it is possible to increase the length of interval *B* without penalizing *ANI*_*c*_. We take advantage of this property to overcome segmentation artifacts, as described later.

Once the matrix of containment indices is computed, ModDotPlot outputs an identity heatmap analogous to a genomic dot plot. The heatmap is assigned a range of color values, ranging from *t* (a user provided threshold identity threshold) to 100. Any cells in the matrix *< t* are left uncolored. Use of *t <* 80 is not recommended, as the identity estimate rapidly loses accuracy below this value for typical values of *k* and *m*, since the higher divergence may result in very few, or zero, *k*-mers shared between the two intervals. Given a symmetric self-identity dot plot, the upper diagonal of the dot plot can be used to produce a triangular dot plot in addition to the standard square.

### Modimizer Hierarchy

Modimizers present a quick and efficient sketching approach, as given a sparsity of *s* = 2^*d*^, only the first *d* bits of each *k*-mer hash need to be checked to verify membership in *MOD*_*s*_. In addition, modimizers are context-independent, providing a guarantee that any *k*-mer selected as a modimizer in one set will also be a modimizer in every other set, regardless of the neighboring context or genomic interval. Given these properties, it is guaranteed that any modimizer in 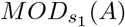 will also occur in 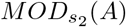 when *s*_1_ is an integer multiple of *s*_2_:

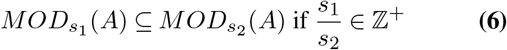

Thus, for a geometric sequence of sparsity values, the smaller modimizer sets will always be subsets of the larger ones. We call this the *hierarchical* property of modimizers. As we describe below, we leverage this property in order to reduce the memory and runtime overhead when generating dot plots at multiple zoom levels.

A *hierarchical modimizer index* consists of *l* modimizer sets with window sizes *ŵ*, 2*ŵ*, …, 2^(*l*−1)^ *ŵ* and corresponding sparsities *ŝ*, 2*ŝ*, …, 2^(*l*−1)^ *ŝ*. Given a user-specified modimizer sketch size 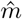 and minimum window size *ŵ*, the initial sparsity is defined as 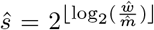. To construct progressively sparser levels of the hierarchy, let *A* be an interval of size 2*w*, and *A*_*L*_ and *A*_*R*_ be the *w*-sized left and right halves of *A* respectively. Due to the hierarchical property, the modimizers for the next sparser level can be sampled from the previous level since *MOD*_2*s*_(*A*) ⊆ *MOD*_*s*_(*A*_*L*_) ∪ *MOD*_*s*_(*A*_*R*_). Repeating this process, additional levels of the hierarchy are sampled until the window size exceeds 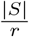, i.e. the resulting number of intervals would be less than the minimum resolution. For example, a 250 Mbp sequence plotted with a minimum window size of 10 Kbp and a resolution of 1,000 would result in 5 layers, since *l* = 5 is the largest *l* such that 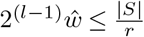. We formally define our algorithm for producing the modimizer hierarchy in Supplementary Algorithm 2.

The runtime and space complexity for building the initial modimizer layer is *𝒪* (*n*), as this requires linear scan of the sequence of size *n*. The expected complexity of each successive layer is half the previous due to the sparsity increasing by powers of two, so the overall runtime and space complexity of Supplementary Algorithm 2 remains *𝒪* (*n*). This approach mirrors the “multilevel winnowing” (12) or “SHIMMER” (7) indices, but our use of modimizers rather than minimizers allows for unbiased containment estimates. From this index, similarity matrices can be efficiently computed for any pair of genomic ranges of the input sequence, with the maximum resolution determined by the minimum window size chosen when building the hierarchy.

### Offset and Window Expansion

When partitioning the input sequence into discrete intervals, it’s possible that two highly similar sequences can be partitioned in different ways, resulting in an inaccurate sequence identity estimate between them (Figure 2). This occurs whenever the two similar sequences are “out of register” and have a different offset relative to the start of the full sequence and that difference is not a multiple of the interval length. The result is that the sequences of the two intervals only partially overlap, rather than fully match. This can also occur within tandem repeats when the unit size is larger than the interval length, such as the rDNA arrays of human acrocentric chromosomes.

**Figure 2.**
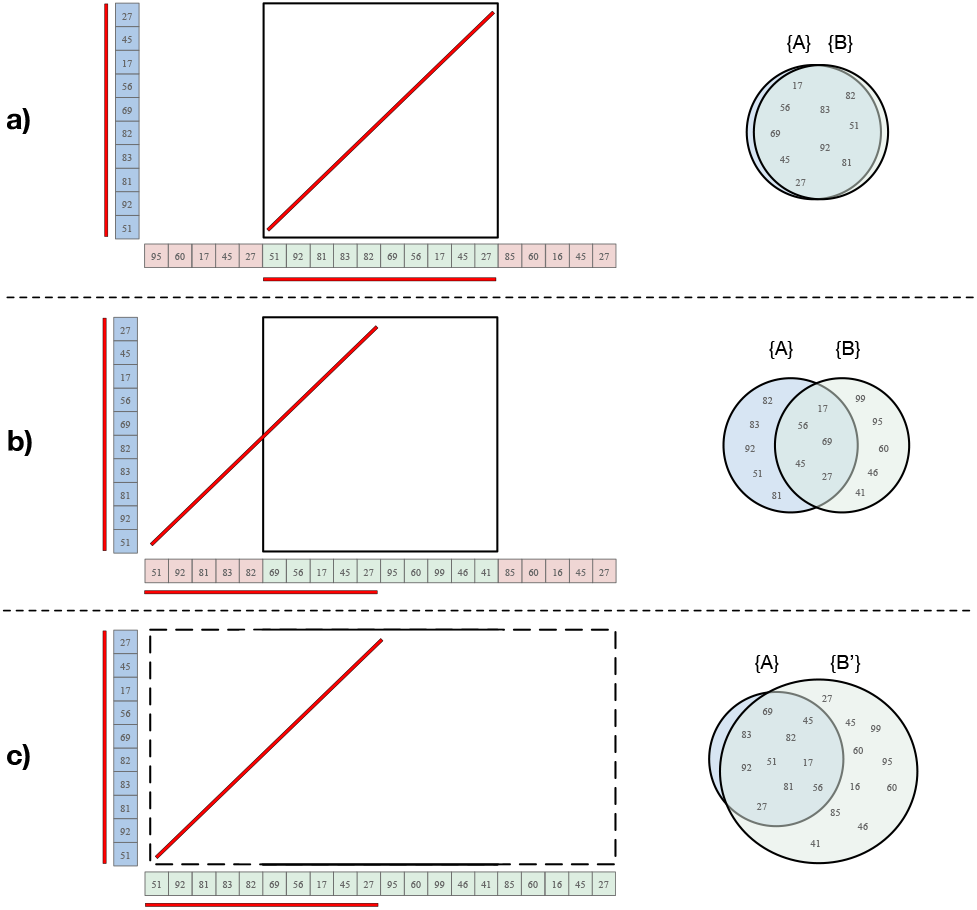
Sample cases for different interval offsets. *k*-mers shared between intervals *A* (blue) and *B* (green) are visualized with a red line. **a)** In an ideal partition, the shared *k*-mers are perfectly captured in both intervals. **b)** In a worse-case partition, only half of the the shared *k*-mers are captured in the cell, leading to a misleading identity estimate for this region. **c)** By keeping *A* fixed, but expanding *B* to *B*^′^, ModDotPlot is able to better capture the similarity between two similar sequences with different offsets. The containment index of *A* in *B*^′^ is then used to determine the score of the dot plot matrix cell *M*_*w*_ (*A, B*).

To overcome this offset issue, ModDotPlot extends each interval *B* by 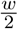 in each direction to form the expanded interval *B*^′^. The containment index is then computed as *c*_*mod*_(*A, B*^′^)^1*/k*^, accounting for any sequence similarities that extend beyond the boundaries of *B*. We show the effect of this approach when computing the containment index in Figure 2, as well as a practical example with human rDNA in Supplementary Figure 1. Since *B* does not appear in the denominator of Equation 4, expanding the size of *B* does not penalize or bias the containment index. Doubling the size of *B* accounts for the worst-case scenario of a match diagonal beginning in the middle of the interval, and so is the default behavior, but this expansion factor can be turned off or adjusted if necessary.

### Implementation and User Interface

ModDotPlot is implemented in the Python programming language (version 3.7 or later). By default, ModDotPlot runs in interactive mode using Plotly (10), which itself uses the Flask web framework (8). Consequently, plots are visualized on a web browser connected to the user’s localhost. Interactive ModDotPlot can also be run remotely, e.g. on a compute cluster, via port forwarding over an ssh tunnel. In static mode, containment indices are saved into a compressed BED file, and dot plots are produced using the Plotnine plotting library (27). In addition to the standard rectangular plots, static mode also supports triangular plot styles.

An important parameter common to all *k*-mer based methods is the choice of *k*, as this represents a trade-off between sensitivity and specificity. Smaller *k*-mers are more sensitive for detecting identity within divergent intervals, but lose specificity due to chance *k*-mer collisions. ModDotPlot allows for flexibility in setting *k*, but based on prior work (25), we set a default *k* = 21 to ensure accurate estimates in most cases.

*k*-mers are hashed using MurmurHash3 (1) and all similarity matrices are stored in the form of NumPy arrays (9). The size of a similarity matrix is proportional to *𝒪* (*r*^2^) rather than the length of the genome sequence. By default, ModDotPlot uses a resolution of *r* = 1, 000 for efficient visualizations on most standard displays. To enable a responsive interface in interactive mode, a full similarity matrix is precomputed for each level of the modimizer hierarchy. However, since the number of layers scales logarithmically with the sequence length, only a few layers are needed in practice (e.g. *l* ≤ 5). When zooming on the plot, the appropriate matrix is chosen such that the number of cells in the matrix is at least the number of pixels in the plot. To prevent redundant computations of similarity matrices for future exploration, NumPy matrices can be saved as binary files and loaded directly as input.

Supplementary Figure 2 shows a screenshot of ModDotPlot’s user interface in interactive mode. Hovering over the plot shows the exact genomic coordinates, along with the corresponding estimated identity of each section. This example shows a plot highlighting the repeat-rich 30 Mbp Y chromosome from a siamang gibbon (*Symphalangus syndactylus*). Users can select a number of preset color-schemes, including high contrast schemes to aid visually impaired or colorblind users, or specify custom colors, either in hex code or RGB format. ModDotPlot also supports the creation of fully-customizable static plots as PDF and PNG files.

## Results

### Plot Accuracy

To showcase the improvements of ModDotPlot over StainedGlass, Figure 3 shows the plots produced by both tools for the centromeric alpha satellite array of the human HG002 X chromosome. The StainedGlass default window size of 2,000 produces a highly “checkered” plot containing streaks of apparently low identity within the array. However, this is not representative of any sort of centromere biology; rather, it is an artifact of partitioning the genome into windows of a fixed size. The canonical DXZ1 higher-order repeat (HOR) present in this array consists of twelve monomers totaling ∼2,050 bp (22), which is slightly longer than the selected window size. Using a window size of 5,000 is sufficient to contain a complete HOR and alleviate this problem, but this comes at the cost of a lower resolution plot and requires advance knowledge of the repeat structure. In contrast, ModDotPlot produces an accurate plot regardless of window length and HOR size.

**Figure 3.**
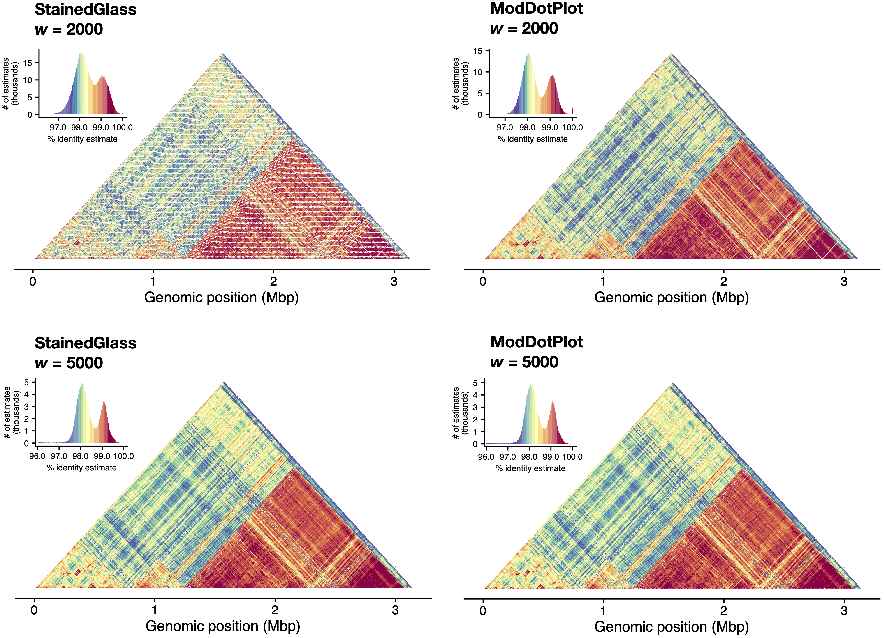
Plots produced by StainedGlass (column 1) and ModDotPlot (column 2), representing the upper diagonal self-identity heatmap of the HG002 DXZ1 satellite array (ChrX:57,680,000–61,000,000). Rows represent a window size of 2, 000 (*r* = 1, 570 in ModDotPlot) and 5, 000 (*r* = 678) respectively. ModDotPlot was run with a default *m* = 1, 000. Plotting artifacts in the StainedGlass *w* = 2, 000 example are due to interactions between the partitioning window size and tandem repeat periodicity.

Figure 4 shows the strong correlation between ModDotPlot *ANI*_*c*_ values and an alignment-based *ANI*_*m*_ computed by MUMmer (21), but with the accuracy of *ANI*_*c*_ decreasing with increasing sparsity (reduced sketch size), as expected (Supplementary Figure 3). For each pairwise combination of HORs present in chrX:58,000,771–58,200,827, the MUM-mer *ANI*_*m*_ was taken from the “AvgIdentity” of 1-to-1 alignments computed by the v4.0.1 “dnadiff” program. The vast majority of HORs, representing the canonical 12-mer structure, fall within the consensus range of 97–100% sequence identity (22) with high concordance (*r* = 0.965) between ModDotPlot and MUMmer. Larger differences between the two methods arise from pairs of windows containing structural variation that confound MUMmer’s alignment-based similarity.

**Figure 4.**
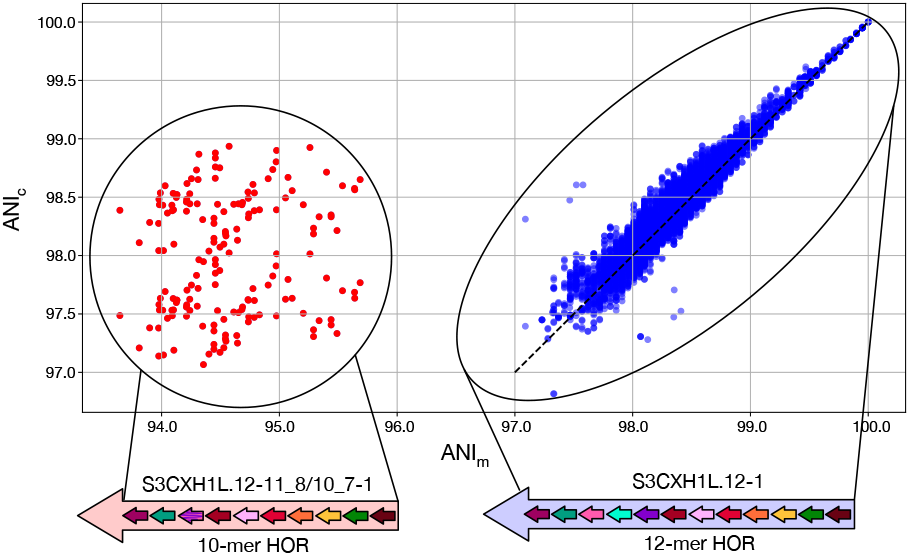
Scatterplots showing the linear relationship between MUMmer *ANI*_*m*_ and ModDotPlot *ANI*_*c*_, using CHM13 chrX:58,000,771–58,200,827. The outlier group labeled in red represents a non-canonical 10-mer HOR (chrX:58,060,405–58,062,120), which is scored differently by the two methods due to the presence of a large deletion when compared to the 12-mer HOR. The dashed line represents *ANI*_*m*_ = *ANI*_*c*_.

The containment index used by ModDotPlot does not penalize *k*-mer copy number differences or large insertions/deletions (indels) in the same way a global alignment would. For example, within the chromosome X centromeric array we observed a small number of windows where the *ANI*_*m*_ and *ANI*_*c*_ values differed substantially. Closer investigation revealed the presence of a single non-canonical HOR, consisting of a shorter 10 monomer repeat that was scored higher by *ANI*_*c*_ when compared to the canonical 12 monomer repeat (Figure 4). The difference between these two repeats is interpreted as a large indel by MUMmer, resulting in a reduced *ANI*_*m*_. However, this difference is not penalized by *ANI*_*c*_, as the 10 monomers present in the shorter HOR are well-contained within the canonical 12 monomer.

Thus, *ANI*_*c*_ is more akin to a local alignment similarity, i.e. the average similarity between the sequences that are shared, and reflects the point mutation rate between two sequences rather than the rate of larger structural variants. This is an important distinction, because in this case MUMmer *ANI*_*m*_ confounds these two evolutionary processes, while *ANI*_*c*_ isolates the point mutation rate of the individual monomers. Such differences between *ANI*_*c*_ and *ANI*_*m*_ are most pronounced within HOR satellite arrays, which are prone to unequal crossing over leading to frequent expansion and contraction of the arrays (3). For this reason, the UniAligner (5) tool, which is specifically built for aligning long tandem repeats, similarly uses an indel penalty of zero during its *k*-mer alignment phase.

### Modimizer Sparsity

Compared to other sketching approaches, modimizers lack any sort of “window guarantee,” meaning that no lower bounds exist on the number of *k*-mers that will be selected for each interval. In addition, the containment index is computed on sets of *k*-mers, not multisets (i.e. only the presence or absence of a *k*-mer is considered), so highly repetitive intervals will typically result in smaller *k*-mer sets, which can lead to reduced accuracy when estimating the containment. Although this is partially taken into account by the error term provided in Equation 4, we demonstrate that by dynamically modifying the sparsity, as done in Supplementary Algorithm 1, the number of modimizers selected per window can be kept above acceptable levels. Figure 5 shows this on a 4 Mbp centromeric region of CHM13 chromosome 1. Regions of alpha satellite repeats show a steep decline in the number of distinct *k*-mers; however, this can be corrected by adaptively reducing the modimizer sparsity in this region to boost the number of *k*-mers selected per window to at least 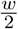 and thus improve the containment estimates. Without this correction, we find that real similarities between low-complexity satellite arrays can go entirely undetected.

**Figure 5.**
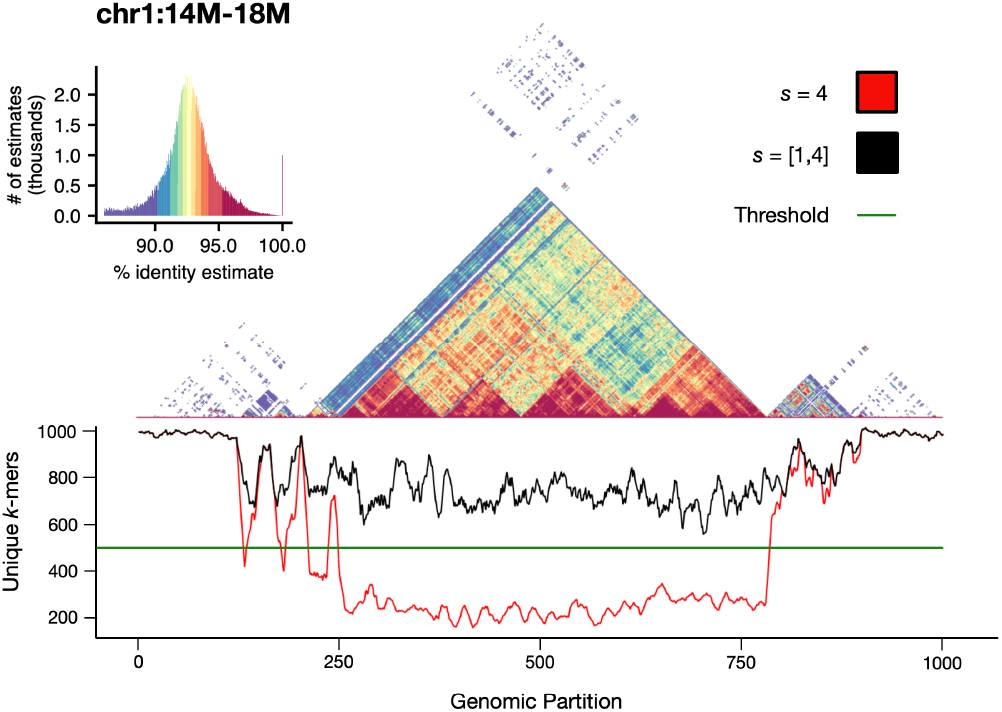
Self-identity plot of the centromere of CHM13 Chromosome 1, overlaid with a smoothed unique *k*-mer frequency chart. Using a window size *w* = 4, 000 and a sparsity *s* = 4, the expected number of modimizers per window is *m* = 1, 000. When using an uncorrected sparsity value (red), the number of unique modimizers per window can drop to under 200. By detecting the unexpectedly small set sizes and adjusting the sparsity of these windows, the total number of modimizers in each window can be increased to at least 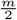 (or, in pathological cases, all *k*-mers in the window).

### Comparative Plots

In addition to self-identity plots, ModDotPlot is also able to generate comparative plots between two different sequences. As an example, we showcase a pairwise dot plot between the DXZ1 alpha satellite arrays of two different human X chromosome centromeres, one from the HG002 genome and one from the CHM13 genome (Figure 6). These two arrays have been previously assembled and compared (3), but it is difficult to understand their structural differences by comparing only their self-identity plots. By plotting the two arrays against each other, their orthology relationship becomes clear. The comparative dot plot of the HG002 and CHM13 DXZ1 arrays reveals a faint diagonal representing the shared history of the two sequences, punctuated by over 300 large duplications/deletions distributed throughout the array (5). As noted above, centromeric satellite arrays are one of the fastest evolving regions of the human genome and accumulate many such structural variants through various recombinational mechanisms. Because of their unique evolutionary patterns, and propensity for bulk insertions/deletions, they have been one of the most difficult regions of the genome to align using traditional approaches.

**Figure 6.**
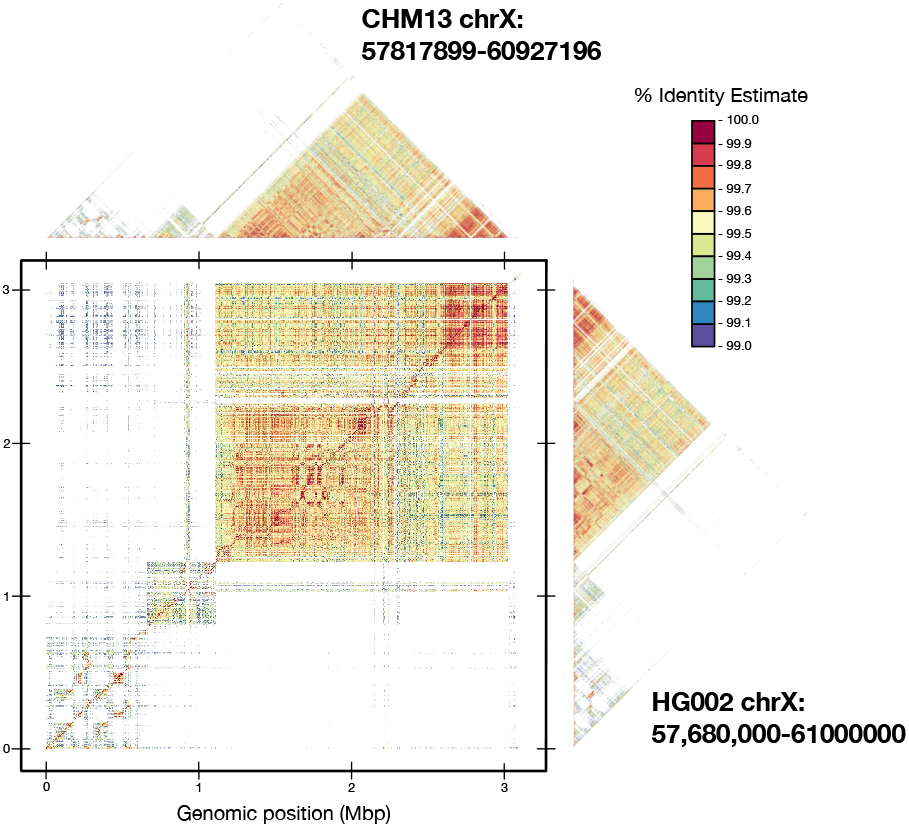
Comparative dot plot of chromosome X DXZ1 satellite array from the HG002 and CHM13 genomes, overlaid with self-identity plots, using a 99% identity threshold. A faint, high-identity diagonal is visible in the comparative plot, indicating the orthologous sequences between these two highly variable arrays.

### Runtime and Memory

n Table 1, we compare the runtime and memory usage of ModDotPlot to StainedGlass across input sequences of various species and sizes. These include the HG002 X chromosome centromere (same sequence as Figure 3), the gibbon Y chromosome (Supplementary Figure 2), the human Y chromosome (30), and the entire gap-free reference genome of *Arabadopsis* (23), containing 5 chromosomes. For each input, both a static matrix and interactive matrices containing three layers were produced, based on a window size proportional to the length of the largest chromosome in the input group. Interactive Stained-Glass plots were created in a similar way to ModDotPlot (i.e. a bottom-up approach based on a minimum window size), and stored in Cooler format (2).

**Table 1.**
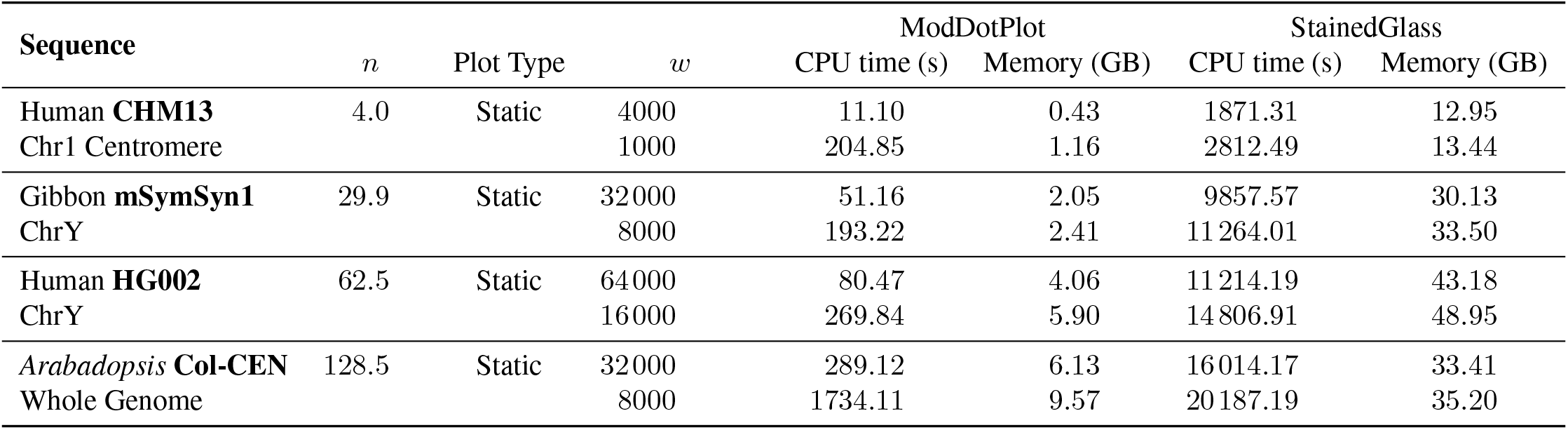
Analysis of memory and runtime needed to produce the similarity matrix (this does not include plot runtime, as that is the same between StainedGlass and ModDotPlot). ModDotPlot was run with a target sketch size of *m* = 1, 000 for all samples. For the *Arabadopsis* Col-CEN assembly, the runtime includes the comparative matrix between each pairwise combination of chromosomes, in addition to self-identity comparisons.

| Sequence | *n* | Plot Type | *w* | ModDotPlot |  | StainedGlass |  |
| --- | --- | --- | --- | --- | --- | --- | --- |
|  |  |  |  | CPU time (s) | Memory (GB) | CPU time (s) | Memory (GB) |
| Human **CHM13** | 4.0 | Static | 4000 | 11.10 | 0.43 | 1871.31 | 12.95 |
| Chr1 Centromere |  |  | 1000 | 204.85 | 1.16 | 2812.49 | 13.44 |
| Gibbon **mSymSyn1** | 29.9 | Static | 32000 | 51.16 | 2.05 | 9857.57 | 30.13 |
| ChrY |  |  | 8000 | 193.22 | 2.41 | 11264.01 | 33.50 |
| Human **HG002** | 62.5 | Static | 64000 | 80.47 | 4.06 | 11214.19 | 43.18 |
| ChrY |  |  | 16000 | 269.84 | 5.90 | 14806.91 | 48.95 |
| *Arabidopsis* **Col-CEN** | 128.5 | Static | 32000 | 289.12 | 6.13 | 16014.17 | 33.41 |
| Whole Genome |  |  | 8000 | 1734.11 | 9.57 | 20187.19 | 35.20 |

In all cases, ModDotPlot exhibits orders of magnitude lower runtime and memory requirements than StainedGlass. An analysis of the Snakemake report generated by StainedGlass showed that the Minimap2 alignment dominated the runtime and memory usage and was the clear bottleneck of the pipeline. We note that despite both tools requiring the sequence identity computation of *r*^2^ cells in each matrix, importantly, ModDotPlot’s runtime is independent of sequence length *n*. Computing *ANI*_*c*_ for each cell requires a set intersection operation on two sets of size *m*, making Equation 5’s runtime complexity *O*(*m*). This can be observed in Table 1, as in interactive mode with high *r*, both Y chromosomes and the Human Chr1 centromere took a similar amount of CPU time, despite each sequence being vastly different in size. In contrast, StainedGlass requires each cell to run Minimap2 on an unsketched sequence of length 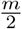. The *𝒪* (*n*) runtime for identity estimation hinders the ability of StainedGlass to visualize whole genomes and large sequences.

## Discussion

Traditional dot plot methods have struggled with the complexity and abundance of genomic repeats, often leading to oversimplified or inaccurate representations. The use of heatmaps offers a substantial improvement over classic vectorized dotplots as they allow for a more natural and nuanced representation of tandem repeats, thereby capturing subtle variations and patterns that vectorized plots obscure. This is especially true for the typical use case where the genomic sequences are manyfold larger than the resolution of the display so that a single pixel intrinsically represents many kilobases of sequence (e.g. a gigabase genome plotted on a 4K display). ModDotPlot improves upon previous methods in terms of speed and computing requirements by an order of magnitude, enabling visualization of whole genomes on a laptop. At the heart of ModDotPlot’s efficiency is its use of hierarchical modimizers, which enable the interactive visualization of vertebrate-sized genomes on a typical laptop. Additionally, the use of expanded intervals combined with the containment index efficiently corrects for registration artifacts inherent to rasterized similarity heatmaps. This is especially important for centromeric and rDNA repeats that are composed of large subunits that can straddle adjacent windows.

A number of additional features could be added to further extend the utility of ModDotPlot. We note how readily satellite arrays and other repeat classes can be visually identified from the dot plots, e.g. satellite arrays appear as dense blocks of color, segmental duplications as lines, and palindromes as lines that cross the diagonal. This raises the possibility of repeat annotation and classification using automated interpretation of dot plots, possibly through machine learning techniques. Additionally, the integration of arbitrary annotation tracks alongside the dot plots would add the ability to visualize genes and other notable features in the context of structural repeats and variation, as is possible with other visualization tools such as HiGlass (13). Lastly, ModDotPlot currently computes similarity matrices in advance of plotting, but with sufficiently fast set operations it would be possible to compute similarity matrices directly from the hierarchical modimizer index on the fly. This would enable interactive exploration of plots with essentially arbitrary resolution.

ModDotPlot highlights the power of minhashing as a fast yet accurate heuristic for sequence alignment, even within the most complex satellite repeat arrays. Alternative sketching approaches may further the utility of this approach. Minmers, for example, allow for an unbiased and accurate identity estimate, with the added advantage of having a window guarantee (14). While such methods can improve sensitivity for more diverged sequences, this comes at the expense of being slower to compute. However, the results presented here suggest that such methods may be able to guide alignments through highly repetitive and variable satellite arrays, ultimately improving our understanding of the structure, function, and evolution of these previously dark regions of the genome.

## Supporting information

Supplementary Algorithm 1, Supplementary Algorithm 2, Supplementary Figure 1, Supplementary Figure 2, Supplementary Figure 3

## Acknowledgements

We would like to thank Mitchell Vollger, Ian Henderson, Karen Miga, and Nicholas Altemose for helpful discussions, and Richard Durbin for suggesting the term “modimizer” to describe an element of a modulo sketch. We would also like to thank Bryce Kille and Nico Ritschel for their feedback and improvements of this manuscript. This work was supported, in part, by the Intramural Research Program of the National Human Genome Research Institute, US National Institutes of Health (A.P.S. and A.M.P.), NSF awards IOS-2216612 and IOS-1758800 (to M.C.S.), and the Human Frontier Science Program award RGP0025/2021 (to M.C.S.). This work utilized the computational resources of the NIH HPC Biowulf cluster (https://hpc.nih.gov).

