## Supplementary Algorithm 1, Supplementary Algorithm 2, Supplementary Figure 1, Supplementary Figure 2, Supplementary Figure 3 for "ModDotPlot—Rapid and interactive visualization of complex repeats"

### Supplemental Material

**Algorithm 1** Partition Sequence S into sets of modimizers

**Input:** List of hashed $k$-mers $S_k[x_1, x_2, ...x_n]$, window size $w$, sparsity $s$
**Output:** List of modimizer sets $MOD_s = [A_1, A_2, ..., A_r]$

**function** BASELAYER($S_k$,$w$,$s$)
  $n \leftarrow$ length of $S_k$
  $r \leftarrow \lceil \frac{n}{w} \rceil$ ▷ Set number of windows (resolution) based on $n$ and $w$
  $m \leftarrow \frac{w}{s}$ ▷ Set the expected modimizer density
  $MOD_s \leftarrow$ list of size $r$
  **for** $i \leftarrow 0$ to $r$ **do** ▷ Retrieve all modimizers within each interval
    $MOD_s[i] \leftarrow \{\}$
    $start \leftarrow wi$ ▷ Set non-overlapping interval boundaries
    $end \leftarrow \min((start + w), n-1)$
    $MOD_s[i] \leftarrow$ GETMODIMIZERS$(S_k[start : end], s, d)$ ▷ Populate list with sets of modimizers
  **end for**
  **return** $MOD_s$
**end function**

**function** GETMODIMIZERS($S_k, s, m$)
  $A \leftarrow \forall x \in \{S_k[A]\} : x \equiv 0 \bmod s$ ▷ Gather the set of unique modimizers per interval
  $\hat{s} \leftarrow \frac{s}{2}$
  **while** $|A| < \frac{m}{2}$ and $\hat{s} > 1$ **do** ▷ Resample at higher density if number of modimizers is under threshold
    $A \leftarrow \forall x \in \{S_k[A]\} : x \equiv 0 \bmod \hat{s}$
    $\hat{s} \leftarrow \frac{\hat{s}}{2}$
  **end while**
  **return** $A$
**end function**

**Algorithm 2** Partition Sequence S into a modimizer hierarchy H

**Input:** List of hashed $k$-mers $S_k[x_1, x_2, ...x_n]$, minimum window size $\hat{w}$, sparsity $\hat{s}$, resolution $r$
**Output:** List $H = [MOD_{\hat{s}}, MOD_{2\hat{s}}, ..., MOD_{2^{l-1}\hat{s}}]$

**function** BUILDHIERARCHY($S_k$,$\hat{w}$,$\hat{s}$,$r$)
  $n \leftarrow$ length of $S_k$
  $l \leftarrow \lfloor \log_2(\frac{n}{\hat{w}r}) \rfloor$ ▷ Initialize number of layers based on min. window size and resolution
  $H \leftarrow$ list of size $l$
  $H[0] \leftarrow$ BASELAYER$(S_k, \hat{w}, \hat{s})$ ▷ Compute bottom layer
  **for** $i \leftarrow 1$ to $l$ **do** ▷ Iteratively compute subsequent layers from previous layer
    $\hat{r} \leftarrow 2^{l-1-i}r$ ▷ Halve the resolution when building subsequent layer
    $H[i] \leftarrow$ ADDLAYER$(H[i-1], 2^i\hat{s}, 2^i\hat{w}, n, \hat{r})$
  **end for**
  **return** $H$
**end function**

**function** ADDLAYER($MOD_{\hat{s}}[A_1, A_2, ..., A_r], s, w, n, \hat{r}$)
  $MOD_s \leftarrow$ list of size $\hat{r}$ ▷ Initialize current layer
  $m \leftarrow \frac{w}{s}$ ▷ Update expected sketch size for current layer
  **for** $i \leftarrow 0$ to $\hat{r}$ **do** ▷ Retrieve modimizers from matching intervals within the previous layer
    $MOD_s[i] =$ GETMODIMIZERS$((MOD_{\hat{s}}[A_{2i}] \cup MOD_{\hat{s}}[A_{2i+1}]), s, m)$
  **end for**
  **return** $MOD_s$
**end function**

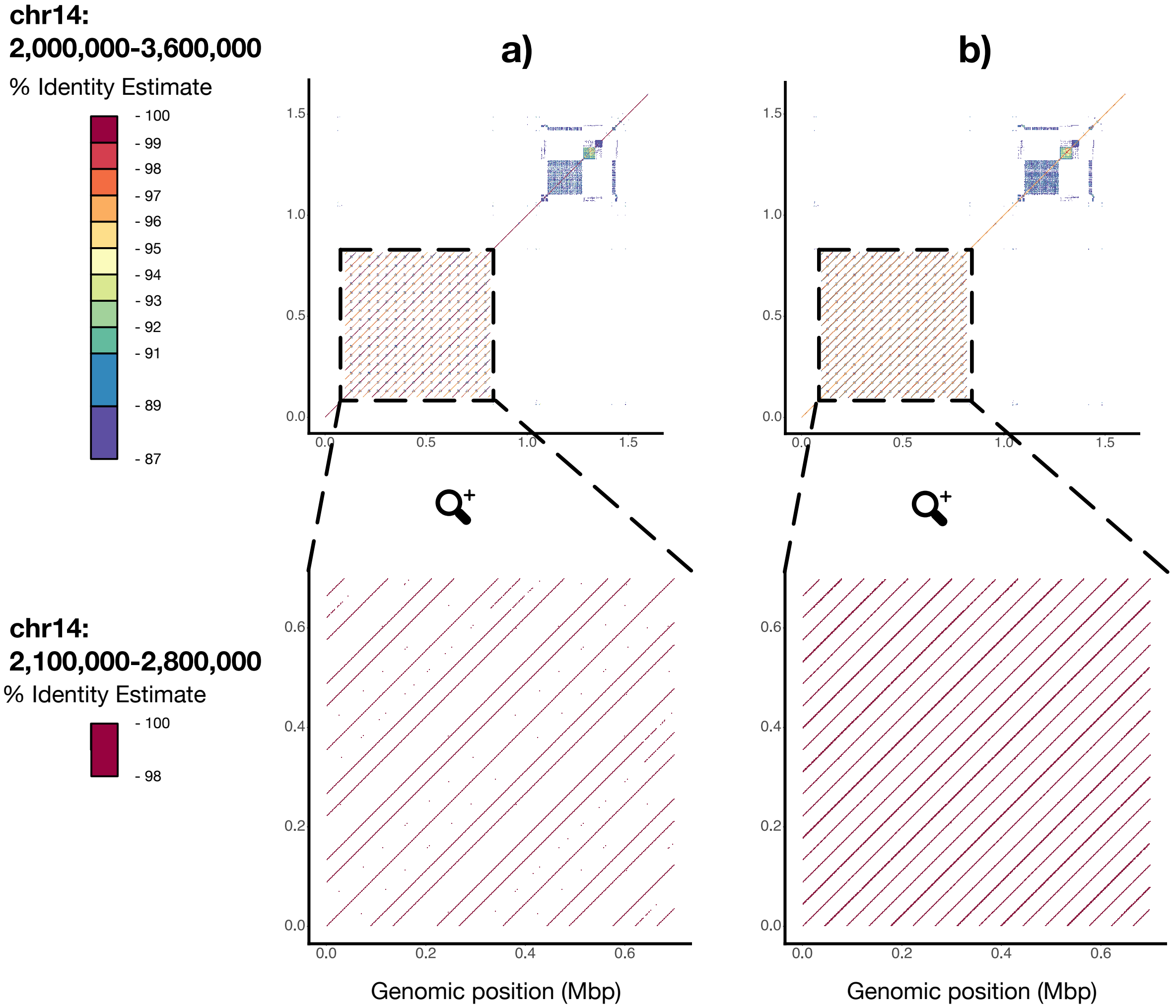


**Supplementary Figure 1.** Screenshots of ModDotPlot run on a human acrocentric short arm (CHM13 chr14:2,000,000-3,600,000), highlighting the rDNA array with and without ModDotPlot's interval extension activated. **a)** With no interval extension, the full 16 copy rDNA array is visible; however, when zoomed in (chr14:2,100,000-2,800,000) and filtered for >98% sequence identity, some rDNA copies disappear from the plot due to registration artifacts. **b)** With intervals extended (i.e. when computing similarity for the cell M(A,B), interval B is extended by $w/2$ in both directions), all rDNA copies appear at all zoom levels.

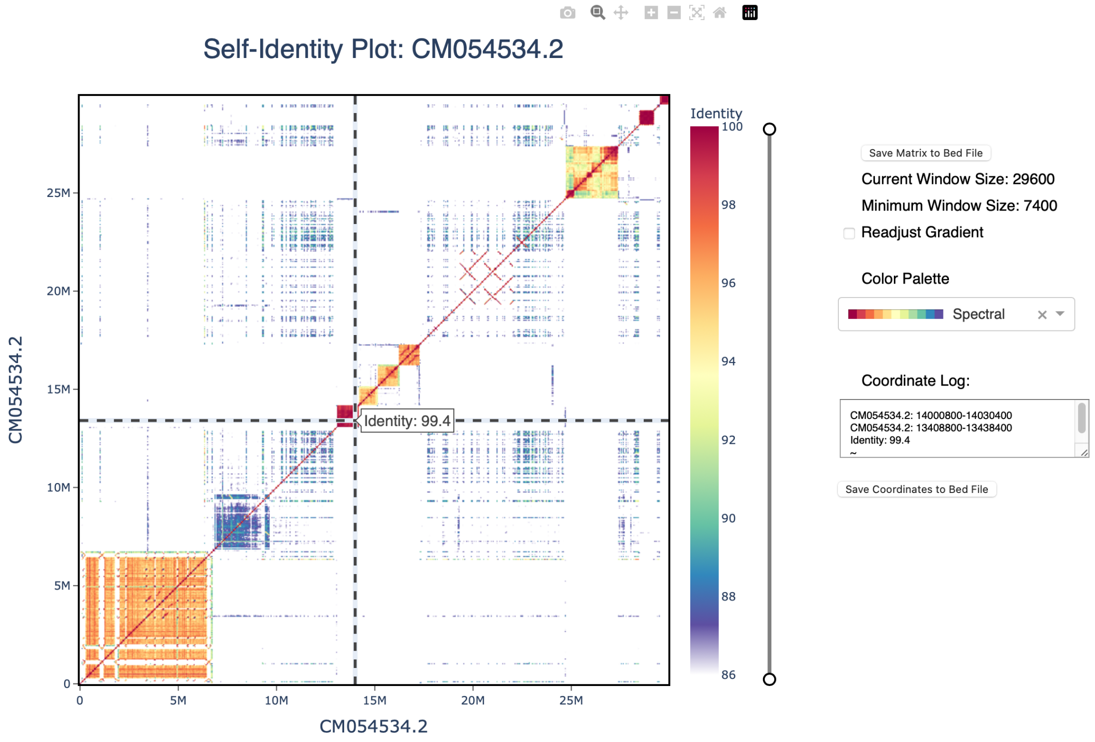


**Supplementary Figure 2.** Screenshot of ModDotPlot's interactive mode, showcasing the entire Y chromosome of a gibbon (mSymSyn1 (1)). Despite spanning almost 30 Mbp, ModDotPlot was able to create 3 matrices in under 2 minutes, with around 2.5GB of memory. Screenshot was taken using ModDotPlot version v0.8.0 (git commit `ed190c7`).

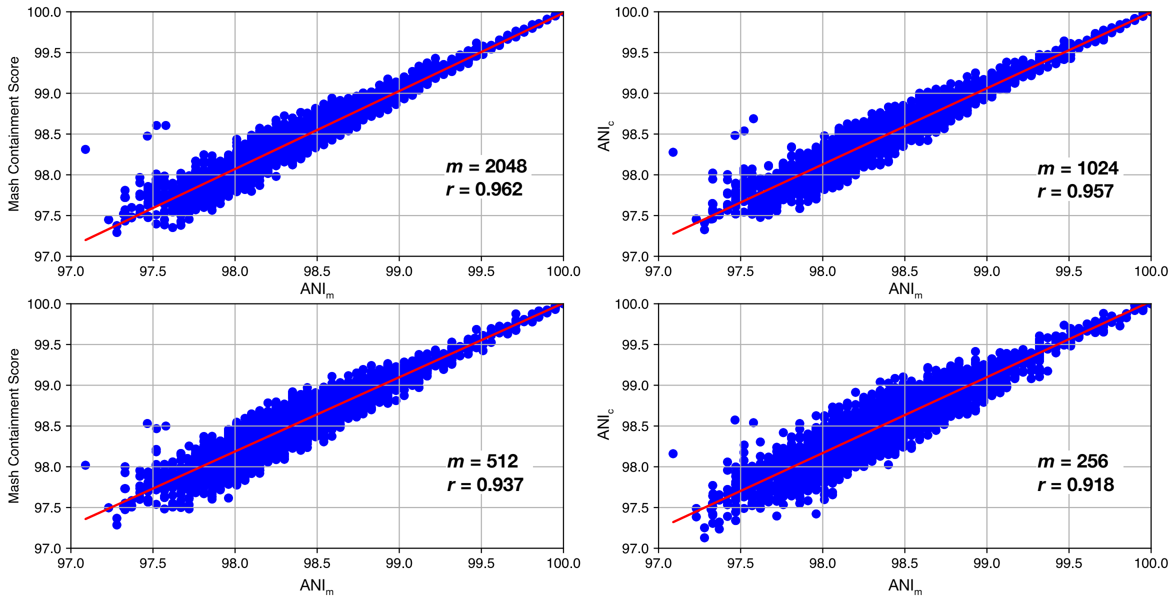


**Supplementary Figure 3.** Scatterplot of $ANI_m$ against the $ANI_c$ distances under various sketch sizes. The 10-mer non-canonical HOR region from Figure 4 was excluded from analysis.

### References

1. Kateryna D. Makova, Brandon D. Pickett, Robert S. Harris, Gabrielle A. Hartley, Monika Cechova, Karol Pal, Sergey Nurk, DongAhn Yoo, Qiuhui Li, Prajna Hebbar, Barbara C. McGrath, Francesca Antonacci, Margaux Aubel, Arjun Biddanda, Matthew Borchers, Erich Bornberg-Bauer, Gerard G. Bouffard, Shelise Y. Brooks, Lucia Carbone, Laura Carrel, Andrew Carroll, Pi-Chuan Chang, Chen-Shan Chin, Daniel E. Cook, Sarah J.C. Craig, Luciana de Gennaro, Mark Diekhans, Amalia Dutra, Gage H. Garcia, Patrick G.S. Grady, Richard E. Green, Diana Haddad, Pille Hallast, William T. Harvey, Glenn Hickey, David A. Hillis, Savannah J. Hoyt, Hyeonsoo Jeong, Kaivan Kamali, Sergei L. Kosakovsky Pond, Troy M. LaPolice, Charles Lee, Alexandra P. Lewis, Yong-Hwee E. Loh, Patrick Masterson, Kelly M. McGarvey, Rajiv C. McCoy, Paul Medvedev, Karen H. Miga, Katherine M. Munson, Evgenia Pak, Benedict Paten, Brendan J. Pinto, Tamara Potapova, Arang Rhie, Joana L. Rocha, Fedor Ryabov, Oliver A. Ryder, Samuel Sacco, Kishwar Shafin, Valery A. Shepelev, Viviane Slon, Steven J. Solar, Jessica M. Storer, Peter H. Sudmant, Sweetalana, Alex Sweeten, Michael G. Tassia, Françoise Thibaud-Nissen, Mario Ventura, Melissa A. Wilson, Alice C. Young, Huiqing Zeng, Xinru Zhang, Zachary A. Szpiech, Christian D. Huber, Jennifer L. Gerton, Soojin V. Yi, Michael C. Schatz, Ivan A. Alexandrov, Sergey Koren, Rachel J. O'Neill, Evan E. Eichler, and Adam M. Phillippy. The complete sequence and comparative analysis of ape sex chromosomes. *bioRxiv*, 2024.
